# Perturbation-aware predictive modeling of RNA splicing using bidirectional transformers

**DOI:** 10.1101/2024.03.20.585793

**Authors:** Colin P McNally, Nour J Abdulhay, Mona Khalaj, Ali Saberi, Balyn W Zaro, Hani Goodarzi, Vijay Ramani

**Author notes:** correspondence may be addressed to, &. these authors contributed equally to this work (Colin P McNally, Nour J Abdulhay, Mona Khalaj).

## Abstract

Predicting molecular function directly from DNA sequence remains a grand challenge in computational and molecular biology. Here, we engineer and train bidirectional transformer models to predict the chemical grammar of alternative human mRNA splicing leveraging the largest perturbative full-length RNA dataset to date. By combining high-throughput single-molecule long-read “chemical transcriptomics” in human cells with transformer models, we train AllSplice – a nucleotide foundation model that achieves state-of-the-art prediction of canonical and noncanonical splice junctions across the human transcriptome. We demonstrate improved performance achieved through incorporation of diverse noncanonical splice sites in its training set that were identified through long-read RNA data. Leveraging chemical perturbations and multiple cell types in the data, we fine-tune AllSplice to train ChemSplice – the first predictive model of sequence-dependent and cell-type specific alternative splicing following programmed cellular perturbation. We anticipate the broad application of AllSplice, ChemSplice, and other models fine-tuned on this foundation to myriad areas of RNA therapeutics development.

## INTRODUCTION

Alternative splicing of messenger RNA (mRNA) is an essential and conserved regulatory process across eukarya^1^. Predicting how DNA sequence encodes alternative splicing programs across different cell-types and cellular perturbations (*i.e*. ‘splicing grammar’) bears promise for both basic scientific discovery (*i.e*. understanding *cis* and *trans* regulators of splicing) and pharmaceutical development (*e.g*. antisense oligo therapies, small-molecule therapy, genome engineering).

The utility of neural networks for this task has become clear with the recent explosion of models capable of predicting how DNA sequence impacts various gene regulatory phenomena, including epigenomic signals^2–4^, chromatin accessibility^5^, transcription factor-DNA interactions^6^, and RNA splicing^7^. While powerful, these applications have been practically limited by the availability of high-information content data. Existing transformer models for predicting RNA splicing from sequence (*e.g*. SpliceBERT^8^) have generally been trained on transcriptomic reference databases such as RefSeq or GENCODE, thus ignoring the large amount of information encoded by cell-type specific RNA processing patterns. Additionally, long-read sequencing has emerged as the gold standard in accurately capturing the complexity of RNA splicing events, yet the current datasets derived from long-read sequencing are relatively small and primarily observational. This highlights a critical need for generating a large, perturbative dataset to harness the full potential of neural networks in understanding RNA splicing dynamics.

To address this limitation, we have developed a complementary approach to understanding splicing grammar. Leveraging a large in-house dataset of high-quality single-molecule RNA sequencing^9^ data from 9 human cell lines treated with 29 different drug treatments, we train AllSplice, a nucleotide BERT model that learns the sequence rules of RNA splicing. Using AllSplice, we demonstrate proof-of-concept for *in silico* saturation mutagenesis, recovering expected rules for canonical splicing, and uncovering novel sequence dependencies for *bona fide* noncanonical splicing events. The perturbative nature of our dataset allows us to go a step further to capture how chemistry perturbs splicing in a context-dependent manner. For this, we present ‘ChemSplice’, a first-in-class fine-tuned derivative of AllSplice that learns cell-type and perturbation-type-specific splicing patterns. We show that ChemSplice predicts splice-site strength in a cell-type specific and drug-treatment specific manner, paving the way for large-scale integration of chemical transcriptomic screens with computational prediction.

## RESULTS

### AllSplice: a splicing-centric RNA foundation model

We sought to train a model capable of predicting where splicing will occur along pre-mRNA sequence. Splice sites (SS) and SS usage are dependent on stereotyped regulatory motifs (*i.e*. splice ‘donor,’ ‘acceptor,’ and ‘branchpoint’ sequences), in addition to other regulatory sequences such as DNA and RNA binding motifs (exonic and intronic splicing enhancers and inhibitors), GC content, and nucleic acid secondary structure. To account for the large variety of potential sequence contributions to SS usage, we leveraged transformer models. “Foundation” transformer models have shown the ability to parse sequence grammars across a variety of domains and have achieved excellent fine-tuning performance for more specialized tasks.

We trained a model (hereafter: ‘AllSplice’) using the BERT architecture, with 12 layers, 12 attention heads, and an internal size of 768 (**Figure 1A**). Unlike existing BERT models, AllSplice is pre-trained on splice junctions themselves, rather than windows of pre-mRNA sequence. Each training example includes windows of sequence (500 basepair [bp] window tokenized as monomers) around each donor and acceptor site. These sequences are tokenized together, analogous to the “two-sentence” input employed by BERT. Uniquely, AllSplice utilizes specialized tokens representing the 5’ and 3’ splice donor and acceptors sites, respectively. AllSplice was pretrained using masked language modeling (predicting whether a splice site is located within a mask). To increase the sensitivity of the model for this task, we increased the assigned weight for masked tokens in the loss calculation, such that predicting masked splice sites was 15% of the loss and predicting masked nucleotides the remaining 85%. Splice site tokens were masked more frequently than nucleotide sequence tokens, to further aid model learning.

**Figure 1.**
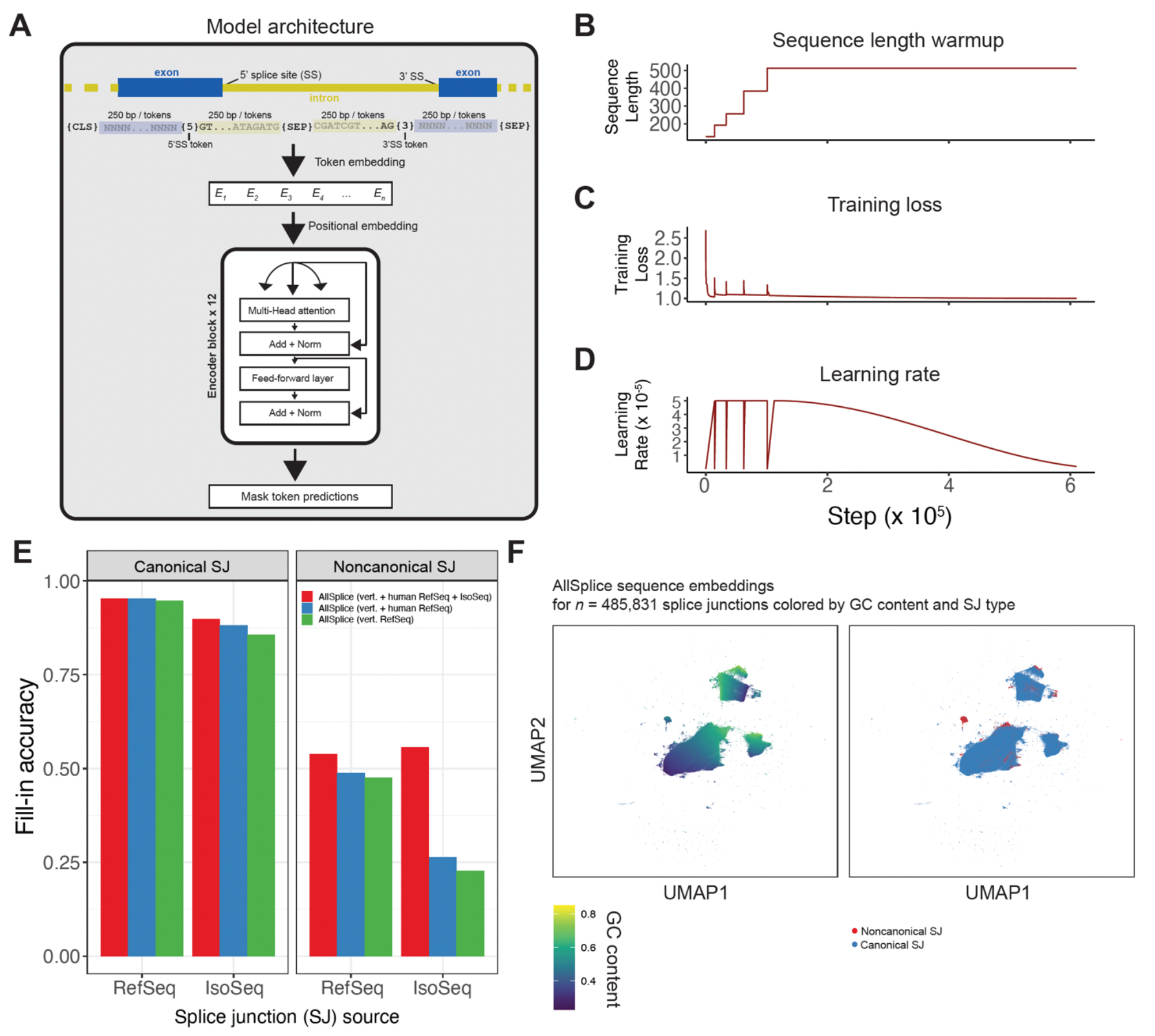
Overview and performance of AllSplice, a model to predict sequence bases for alternative splicing of mRNA. **A.)** Schematic model of AllSplice, a transformer model trained on the RefSeq database and single-molecule long-read “IsoSeq” data. **B.)** AllSplice was trained using sequence length ‘warmup.’ **C.)** Model training loss as a function of training step. **D.)** Model learning rate as a function of training step. **E.)** AllSplice performance, stratified by training corpus and by splice junction (SJ) source. F.) UMAP visualization of AllSplice sequence embeddings for 485,831 SJs, colored by GC content (**left**) and canonical vs. noncanonical SJ type (**right**).

The final AllSplice model comprises an initial model pre-trained on a large set of splice junctions from RefSeq, which was then further trained using proprietary singe-molecule IsoSeq data generated by Entwine Bio. We initially pre-trained the model on junctions taken from all vertebrate transcriptomes, filtered for mRNA transcripts whose model evidence included 100% RNA coverage and at least one sample with complete coverage (in total, 4.3E7 splice junctions from 326 species). After training this initial model, AllSplice was then fine-tuned on human-specific splice junctions only – 229,163 derived from RefSeq, and 177,141 junctions derived from Entwine’s dataset. All IsoSeq junctions were filtered by requiring supporting short-read coverage for the same junctions, at least two occurrences of the junction in the IsoSeq dataset, and requiring that all introns were > 50 bp long.

AllSplice was trained using sequence length warmup^10^ (**Figure 1B**), gradually expanding the size of nucleotide windows around each splice site. The final sequence length was 1024 tokens; the initial model was trained for 6E5 steps, and the final model was fine-tuned for 7E4 steps using human-specific / IsoSeq data (**Figure 1C,D**). We evaluated model performance by computing prediction accuracy for splice site placement when tokens were masked (*i.e*. ‘fill-in accuracy’; performance for models trained on RefSeq only [RO], RefSeq plus fine-tuning on human junctions [RH], and all data [AllSplice] tabulated in **Figure 1E**). For held-out human RefSeq data spanning canonical SJs, AllSplice achieved fill-in accuracy of 95.3% (compared to RO: 94.7%; RH: 95.4%). For held-out IsoSeq specific canonical SJ, AllSplice achieved accuracy of 89.7% (RO: 85.6%; RH: 88.1%). For held-out human RefSeq data spanning noncanonical SJs, AllSplice achieved accuracy of 53.7% (RO: 47.5%; RH: 48.8%). For held-out IsoSeq noncanonical SJ, AllSplice achieved accuracy of 55.6% (RO: 22.7%; RH: 26.4%).

We next visualized the learned sequence embeddings from AllSplice using the UMAP algorithm^11^. Positions of pre-mRNA sequences in the *k*-nearest-neighbor graph UMAP representation broadly tracked with sequence features known to impact splicing, including GC content (**Figure 1F;** left), and the identity of a given splice junction (SJ; *i.e*. canonical vs. noncanonical; **Figure 1F;** right). Together, these analyses demonstrate the ability of AllSplice to learn splicing grammar for canonical and noncanonical SJs specific to long-read-based transcriptomes.

### AllSplice enables *in silico* saturation mutagenesis of canonical splice sites

A common application of nucleotide models is *in silico* saturation mutagenesis (ISM), which enables rapid variant effect prediction. We sought to determine if AllSplice could be applied to a similar task to estimate impacts of single-nucleotide variants (SNVs) on splice junction formation, integrating information from both 5’ and 3’ SS. We devised an ISM approach that works as follows (schematized in **Figure 2A**): i.) we isolated 1,290 canonical splice junctions to mutagenize, extracting 500 nt of genomic sequence around each 5’/3’ SS; ii.) to account for outsized effects of individual masked nucleotides, for each junction we mutagenize 10 masked test sequences, each with a random 20% of nucleotides masked; iii.) we systematically mutagenized and tested every possible mutation for every junction across all masked starting sequences; and iv.) for each test, we recorded both the mutation, and the predicted effect of each mutation. Mutation effects were quantified as “Δ”, a representation of the change in AllSplice’s probability estimate for the true splice site (effectively, an estimate of change in splice site strength). These values were calculated for both nearby SS (*i.e*. ‘proximal’ effects) and the matched splice donor / acceptor (*i.e*. ‘distal’ effects).

**Figure 2:**
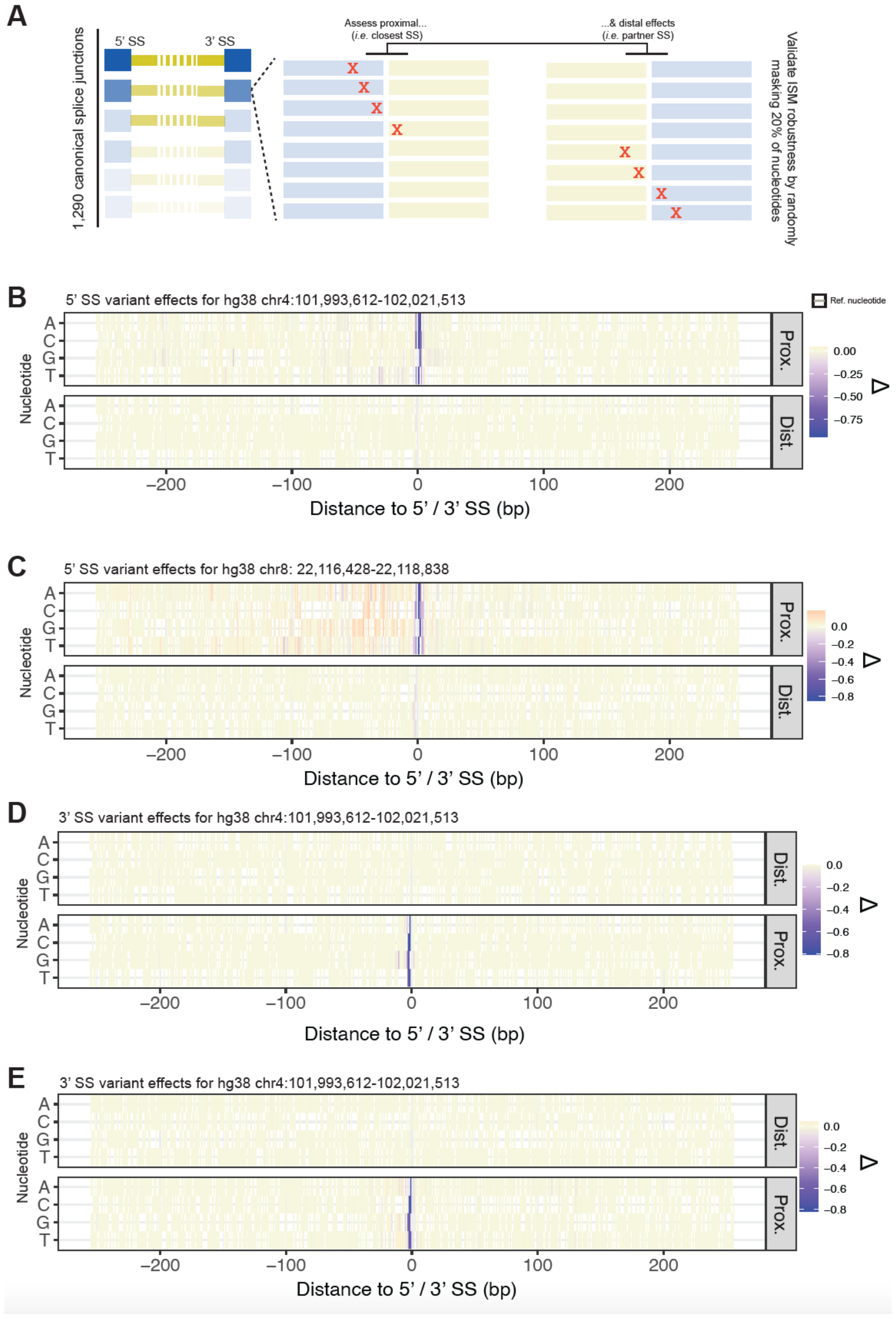
AllSplice enables *in silico* saturation mutagenesis of canonical splice junction for variant effect prediction. **A.)** Schematic of *in silico* saturation mutagenesis (ISM) approach using the AllSplice model. For each of 1,290 canonical SJs, we tested how single nucleotide changes within a 500 bp window surrounding either the 5’SS or 3’SS impact the probability of detecting the expected site. ISM was carried out on templates that with 20% of nucleotides randomly and iteratively masked to ensure robustness of ISM measurements. **B.-E)** Heatmap representations of ISM scores (*i.e. ‘*Δ’), representing the change in the probability that the predicted splice site is correctly positioned, for two different splice sites found in the human reference genome. (**B-C**) represent effects for the 5’SS for both loci, and (**D-E**) represent effects for the 3’SS for both loci.

We computed the effects of mutations for all possible mutations across 485,831 human splice sites seen by AllSplice (**Figure 2B-E**). For two of these splice sites, we visualized these effects as quartets of heatmaps, corresponding to the proximal and distal predicted variant effects for each non-reference nucleotide at 5’ and 3’ SS. Predicted variant effects localized to the vicinity of the splice donor and acceptors themselves, with muted— though nonzero—predicted distal effects for both visualized SJs. Intriguingly, AllSplice predictions were junction-specific (compare **Figure 2B,D** to **Figure 2C,E**), with SJs demonstrating unique sequence dependencies outside of the canonical donor and acceptor sites, such as predicted exonic effects on splice site strength. Together these analyses demonstrate the capability of AllSplice for nominating sequence variants that directly impact splice donor and acceptor choice.

### AllSplice predicts novel sequence dependencies at noncanonical splice sites

A unique advantage of AllSplice compared to existing RNA language models is the inclusion of noncanonical splice sites discovered via IsoSeq in its training corpus. We thus sought to quantify AllSplice’s learned sequence dependencies at canonical versus noncanonical splice sites via ISM. We first visualized the magnitude of mutation effects for both classes of sites as distributions of predicted effects for all nucleotides, plotted with respect to distance from 5’ / 3’ SS. While canonical splice sites (**Figure 3A**) were predicted to be highly impacted by mutations to the splice donor and acceptor sites, we observed a much broader range of effects to nucleotides in the vicinity of noncanonical splice sites (**Figure 3B**). The variability of nucleotide effects, and the distance over these which these effects were exerted is illustrated in an example ISM for a noncanonical SJ found in the *ZNG1E* gene (**Figure 3C,D**). Zoomed-in visualization of the ISM heatmap for this SJ reveals a wide range of predicted variant effects, including ‘distal’ variants occurring near the 3’SS with weak positive effects on the 5’SS (**Figure 3C**), and both positive and negatively acting proximal 3’SS variants (**Figure 3D**). These analyses demonstrate proof-of-concept for future studies leveraging AllSplice to dissect transcriptome-scale rules for noncanonical SJ usage.

**Figure 3.**
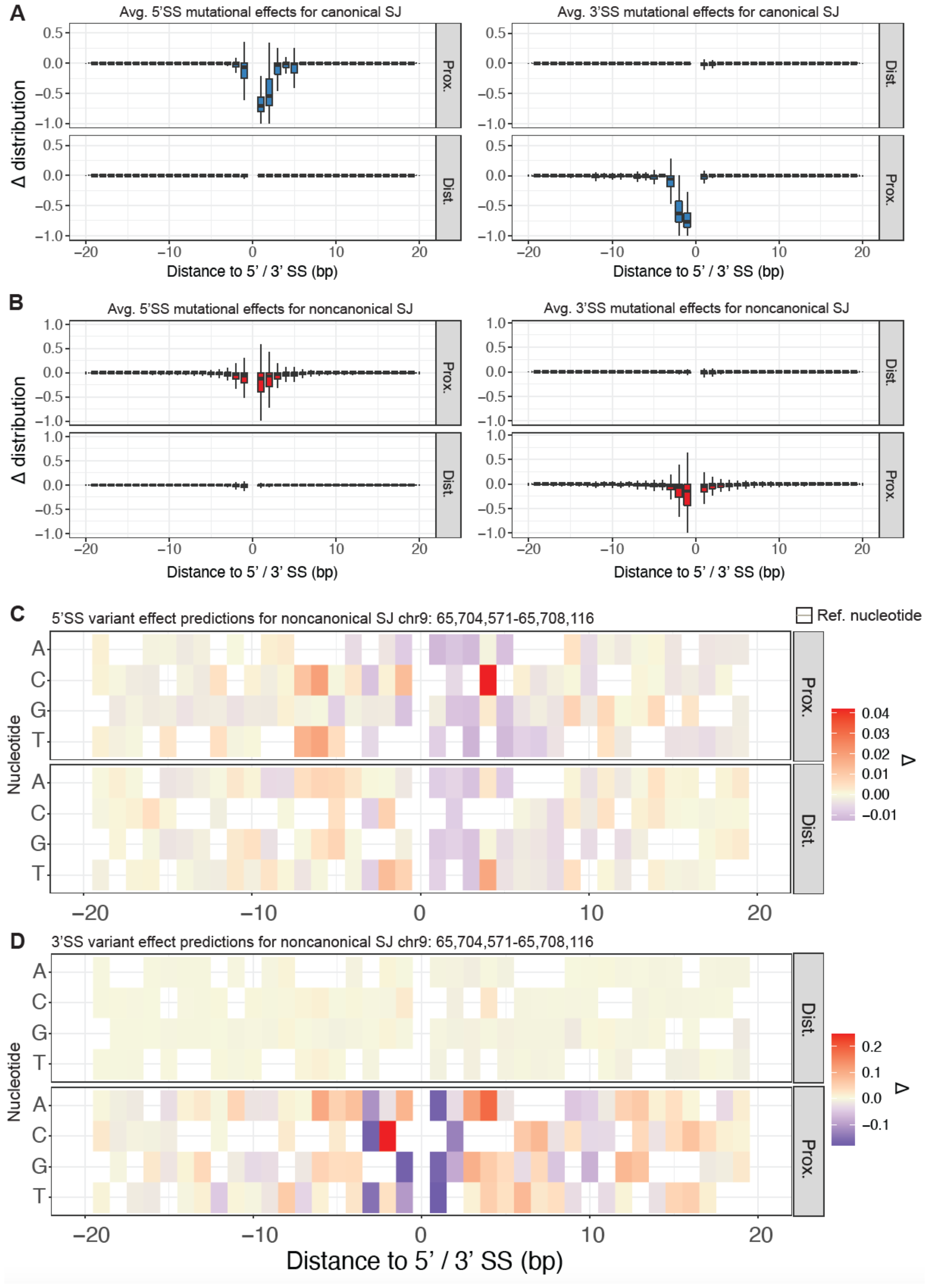
Comparison of model predictions and ISM of canonical versus noncanonical splice junctions. **A.)** Boxplot representation of the average and distribution of predicted variant effects for (**A**) canonical 5’ and 3’SS and (**B**) noncanonical 5’ and 3’SS. **C-D.)** Heatmap representations (zoomed in compared to **Figure 2**) of variant effect predictions for 5’ SS (**C**) and 3’SS (**D**) for a SJ spanning locus chr9: 65,704,571 – 65,708,116.

### ChemSplice learns cell-type and drug-perturbed splicing grammar

Existing nucleotide transformer models are agnostic to cellular state and cellular perturbations, omitting major axes of biologically-relevant variation. To go beyond sequence-only predictions, we developed a model termed ‘ChemSplice,’ which attempts to build an explicit contextual understanding of how cell-type specificity and drug perturbations interact to alter splicing grammar. The model architecture for ChemSplice is schematized in **Figure 4A**. ChemSplice utilizes three primary sources of input: sequence embeddings for splice sites provided by AllSplice, cell-type specific gene expression embeddings (generated via principal components analysis), and drug embeddings provided by ChemBERTa-2^12^ activation and dropout. The output of these layers uses a sigmoid activation function to yield a probability between 0 and 1, which can be interpreted as individual splice-site strength (SSE)^13^.

**Figure 4:**
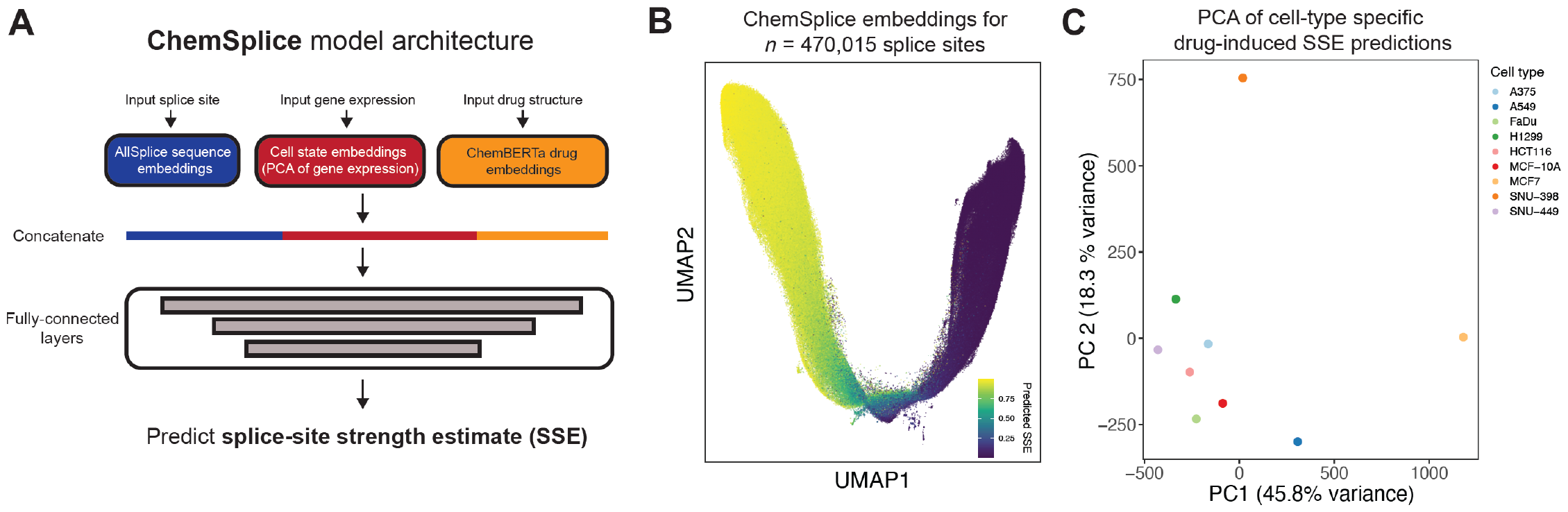
ChemSplice integrates cell-type specific gene expression measurements and drug-perturbation data to capture perturbation-specific splicing grammar. **A.)** Schematic of the model architecture for ChemSplice. The model integrates three separate embeddings: splice site embeddings provided by AllSplice, gene expression embeddings generated by principal components analysis of cell-type specific gene expression data, and chemical compound structural embeddings provided by ChemBERTa-2. These embeddings are concatenated and fed into fully-connected layers to yield a value from 0 – 1 representative of a splice-site strength estimate. **B.)** UMAP visualization of sequence embeddings for 470,015 splice sites seen by ChemSplice, colored by mean predicted SSE. **C.)** PCA-based reduced dimension visualization of a cell type x splice-site matrix, where matrix elements are the predicted variance in SSE across all drug treatments for a given cell type. This visualization indicates that ChemSplice is capturing cell- and drug-perturbation-specific information in its predictions.

ChemSplice’s loss function is the log-likelihood of the number of observed reads utilizing the splice site, given the splice site usage probability output by the model. The likelihood was calculated from the binomial function. This loss function accounts for uncertainty in the measurements of splice site usage from RNA-seq data, by weighing more highly the sites with larger numbers of reads that thus have a more accurate estimate of the fractional site usage. ChemSplice was trained for 35 epochs, using 8X A100 40GB GPUs. The final two encoder layers of Allsplice were removed due to being specialized for the masked language modeling task.

We first examined the sequence embeddings generated by ChemSplice. Sequences were broadly arranged in *k*-nearest-neighbors space according to the mean SSE for each splice site across cell lines and drug treatments (**Figure 4B**), indicating the ChemSplice has learned relationships between splice site sequences and splice site usage from the data. We next sought to determine if ChemSplice captured cell-type and drug-type specific variation. To test this, we examined the variance in SSE for each splice site, for each cell type tested, across all drugs tested, and then visualized the relationship between cell types on the basis of drug-induced SSE variance at each splice site using PCA (**Figure 4C**). Intriguingly, cell types clustered into distinct regions in PCA space, suggesting that ChemSplice has captured aspects of cell-type and drug-perturbation specific changes in splice-site strength. Taken together, these analyses demonstrate the ability to train models capable of integrating splice site sequence information, cell-type specific expression information, and encoded perturbation information to learn condition-specific aspects of splicing grammar.

## DISCUSSION

Predictive models capable of understanding the consequence of biological perturbation remain difficult to engineer. Towards addressing this challenge, we present two related models, AllSplice and ChemSplice; together these comprise novel predictive tools for understanding how DNA sequence, cell state, and cellular perturbations impact alternative splicing of human mRNA. AllSplice is a BERT model that contains splicing as an explicitly defined component of its input and output, trained on reference transcriptomes and long-read RNA sequencing data. We leverage this unique advantage of AllSplice to demonstrate proof-of-concept for variant effect prediction at both canonical and noncanonical splice junctions across the human transcriptome. ChemSplice combines AllSplice embeddings, cellular gene expression, and chemical entities to predict perturbed splice site usage; we demonstrate that ChemSplice captures aspects of cell-type specific RNA splicing grammar.

Most existing sequence based foundation models have aimed to learn fundamental DNA and RNA biology through pre-training on sequence alone. We hope to demonstrate the added potential of customizing the model architecture and data input to achieve specific tasks that require an increased level of understanding of the interaction that nucleotide sequence has with cellular and environmental factors. Further, predictive technologies are only as good as the molecular experiments they inspire. In future work, we anticipate combining AllSplice and ChemSplice predictions with high-throughput functional validation, which can then be used to further fine-tune the model. Combining such data with an ever-growing data corpus of diverse perturbations measured by high-quality long-read transcriptomics holds much promise for both basic and translational RNA biology.

